# SomaticSiMu: A mutational signature simulator

**DOI:** 10.1101/2021.09.30.462618

**Authors:** David Chen, Gurjit S. Randhawa, Maximillian P.M. Soltysiak, Camila P.E. de Souza, Lila Kari, Shiva M. Singh, Kathleen A. Hill

## Abstract

**Summary:** SomaticSiMu is an *in silico* simulator of single and double base substitutions, and single base insertions and deletions in an input genomic sequence to mimic mutational signatures. SomaticSiMu outputs simulated DNA sequences and mutational catalogues with imposed mutational signatures. The tool is the first mutational signature simulator featuring a graphical user interface, control of mutation rates, and built-in visualization tools of the simulated mutations. Simulated datasets are useful as a ground truth to test the accuracy and sensitivity of DNA sequence classification tools and mutational signature extraction tools under different experimental scenarios. The reliability of SomaticSiMu was affirmed by 1) supervised machine learning classification of simulated sequences with different mutation types and burdens, and 2) mutational signature extraction from simulated mutational catalogs.

**Availability and Implementation:** SomaticSiMu is written in Python 3.8.3. The open-source code, documentation, and tutorials are available at https://github.com/HillLab/SomaticSiMu under the terms of the Creative Commons Attribution 4.0 International License.

**Supplementary information:** Supplementary data are available at *Bioinformatics* online.

## 1 Introduction

Somatic mutagenesis in cancer arises from the interplay between inherited genomic instabilities, acquired mutations associated with error-prone replication and faulty repair, as well as endogenous and exogenous DNA damaging mechanisms. Sources of somatic mutagenesis are known to generate characteristic patterns of small-scale base substitutions, insertions, and deletions known as mutational signatures that cumulatively alter cancer genome sequence composition (Supplementary Data A) (Alexandrov *et al*., 2020; Bacolla *et al*., 2014; Forbes *et al*., 2016).

Optimizing sequence classification tools and mutational signature extraction tools can be achieved by systematically assessing their performance using simulated datasets. First, mutation simulations offer the capacity to generate large datasets as input to benchmark machine learning-based tools for supervised classification of genomic sequences with specific predefined mutational signatures. Simulated sequences can be used in the case of rare cancer genomic sequences to upsample minority classes to correct systematic performance biases by machine learning-based sequence classifiers against minority classes. Second, ground truth genomic sequence datasets with imposed mutational signatures and known sets of mutation types, contexts, and burdens can also be used to benchmark the accuracy of mutational signature extraction tools. SomaticSiMu is a new mutational signature simulator with confirmed utility for these applications (Supplementary Section B).

While genomic sequences from cancer biosamples remain the gold standard input for benchmarking the performance of machine-learning based sequence classification tools, there remain several advantages of simulating genomic sequences *in silico* to complement validation with real data (Mangul *et al*., 2019). Simulation allows for the fine-tuned control of parameters needed to generate large and highly customized sequence datasets. Moreover, simulated sequences have known sets of mutation types and burdens while real sequences are susceptible to random error in mutation detection due to sequencing artifacts, incorrect local read alignments, and tumor heterogeneity (Semeraro *et al*., 2018).

Current tools used to simulate mutations associated with cancer include SigProfilerSimulator (Bergstrom *et al*., 2020), Xome-Blender (Semeraro *et al*., 2018), esiCancer (Minussi *et al*., 2018), simuG (Yue *et al*., 2019), and Simulome (Price *et al*., 2017). Compared to SomaticSiMu, these simulators do not simulate mutations on custom subsets of the genome (e.g., SimuG), do not output DNA sequence FASTA files needed to evaluate the performance of sequence classification tools (e.g., esiCancer and SigProfilerSimulator), or are not designed to simulate the mutational types and burdens associated with multiple mutational signatures at once (e.g., Xome Blender and Simulome).

## 2 Features

SomaticSiMu is a standalone software tool that simulates mutational signatures operative in human cancer genomes. SomaticSiMu offers a range of features that allows users to simulate realistic mutation types and proportions associated with known single base substitution (SBS) and double base substitution (DBS) mutational signatures, and single base insertions and deletions (indel) from the Catalogue of Somatic Mutations in Cancer (COSMIC) database (Tate *et al*., 2018). Each simulation is designed to mimic the multiple mutational signatures observed in the user-selected human cancer type, each associated with different burdens and attributed to a source of mutagenesis. SomaticSiMu offers a simulation mode to generate simulated sequences with imposed mutational signatures (Supplementary Section C) and a visualization mode to plot mutational signature plots and total mutational burden for each simulated sequence (Supplementary Section D), see Figure 1 for an overview of the SomaticSiMu pipeline.

**Fig. 1.**
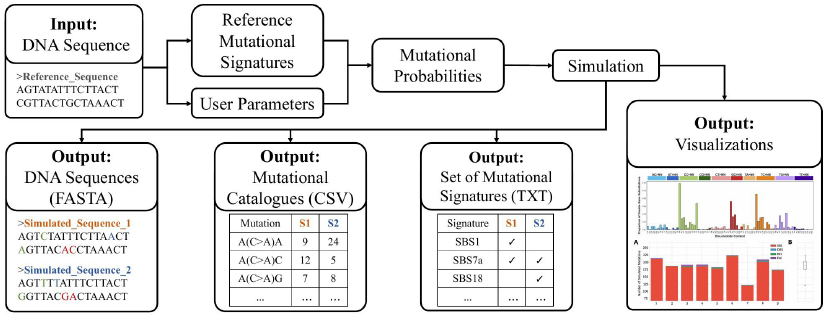
SomaticSiMu Pipeline. SomaticSiMu pipeline structure and simulation features.

## 3 Methods

SomaticSiMu first preprocesses the input reference sequence by mapping the sequence *k*-mer composition. Next, SomaticSiMu samples the set of SBS, DBS, and single base indel signatures operative in a user-specified cancer type. The tool samples subsets of active signatures based on their prevalence among COSMIC tumor samples associated with the user-specified cancer type for each simulation. Mutation probability is defined as the probability that any given mutation type found in the SBS-96, DBS-78, and single base indel classification schemes will occur at a specific local sequence context type in the simulated genome. SomaticSiMu introduces novel mutations in the reference sequence at random indexes, based on the vector of mutation types and their associated mutation probabilities derived from reference mutational signatures and user-defined controls.

## 4 Performance Evaluation

To assess the speed and memory consumption of SomaticSiMu, we simulated 20 genomic sequences, each for five cancer types, using the 50 Mb Chromosome 22 from the human genome assembly GRCh38.p13 (NCBI accession: NC_000022.11) as the input reference sequence (Supplementary Table S4). Performance tests were conducted on a MacBook Pro *A*2141 with a 2.3GHz 8-core Intel Core *i*9 9880H processor, 16GB DDR4 2667MHz SDRAM, and eight parallel processes. SomaticSiMu took less than 700 seconds to complete each simulation run and recorded a peak memory consumption of less than 10GB.

Eleven benchmark experiments (Supplementary Section E) suggest that SomaticSiMu can accurately simulate mutation types and proportions comparable to mutation types and proportions observed in cancer genome sequences from COSMIC (cosine similarities > 0.95, and 11 out of 12 Wilcoxon rank-sum tests indicating no significant difference, *p* > 0.05).

To test the reliability of SomaticSiMu, sequences with combinations of high or low mutational burden and high or low cosine similarity between mutational signatures were simulated. A sequence classification tool (Machine Learning with Digital Signal Processing, MLDSP; Randhawa *et al*., 2019) was used to confirm the expected nature of the simulations. Five experimental tests showed that simulated sequences with higher total sequence mutational burden and more dissimilar mutation types faithfully reproduced the expected increase in MLDSP classification accuracy, from a minimum accuracy of 34.8% up to a maximum accuracy of 72.4% using the best-performing Linear SVM classifier (Supplementary Section F). Also, simulated mutational catalogues with high and low mutation counts were used to extract mutational signatures. Signature extraction tested on 100 simulated mutational catalogues showed that SigProfilerExtractor (Islam *et al*., 2021) consistently extracted the subset of simulated mutational signatures with high mutation counts as expected, up to a maximum specificity of 100% and sensitivity of 79.5% (Supplementary Section G). Both the classification of simulated sequences and the extraction of mutational signatures from simulated mutational catalogues affirmed the accurate simulation of mutational signatures.

## Acknowledgements

We thank Akshaj Jonnalagadda and Adrian Yung for testing SomaticSiMu.

