## Supplementary for "SomaticSiMu: A mutational signature simulator"

### SomaticSiMu Documentation
### Supplementary Material

D. Chen[1], G. S. Randhawa[2], M. P. M. Soltysiak[1], C. P. E. de Souza[3],
L. Kari[2], S. M. Singh [1], K. A. Hill[1].
[1]Department of Biology, Western University, London, Ontario, Canada
[2]School of Computer Science, University of Waterloo, Waterloo, Ontario, Canada.
[3]Department of Statistical and Actuarial Sciences, Western University, London, ON, Canada

#### A Overview of Mutational Signatures

Somatic mutations associated with cancer can be caused by a variety of exogenous and endogenous sources of mutagenesis. Since the advent of next-generation sequencing technologies, it is possible to identify a catalogue of the thousands of mutations from tumor genomes that result from the interplay of multiple mutational processes. Each mutational process may be associated with one or more mutational signatures. A mutational signature is defined as the characteristic combination of mutation types, mutation local sequence contexts, and mutation proportions attributed to a mutational process (Alexandrov *et al*., 2020). Multiple mutational processes can be active at once in one tumor, leading to the observation of multiple active mutational signatures. The activity of multiple mutational processes can lead to multiple observed mutational signatures that collectively make up the total catalogue of mutations and mutational burden observed in a tumor genome.

The dominant classes of substitution mutational signatures can include single base substitution (SBS) signatures and double base substitution (DBS) signatures. The reference base of a substitution is identified by the pyrimidine of the Watson-Crick base pair (Alexandrov *et al*., 2020). The 96-type SBS mutational signature (SBS-96) refers to the 6 SBS types in the local trinucleotide context (see Figure S1A for an example SBS-96 visualization and Figure S1B for the breakdown of the SBS-96 classification scheme). The 78-type DBS mutational signature (DBS-78) refers to 10 strand-agnostic DBS types (see Figure S1C for an example DBS-78 visualization and Figure S1D for the breakdown of the DBS-78 classification scheme).

Single base insertions and deletions (indels) describe the incorporation or loss of a single base at a specific site in the genome respectively. Due to the difficulty in defining a consistent set of indel mutation types, one approach used to describe the set of single-base indels is to consider their occurrence at mononucleotide repeat regions with a length of 1 base up to 6 or more bases. The classification of single base indels at repeat regions refers to 12 insertion types and 12 deletion types (see Figure S1E for an example indel visualization and Figure S1F for the breakdown of the 12-type indel classification scheme).

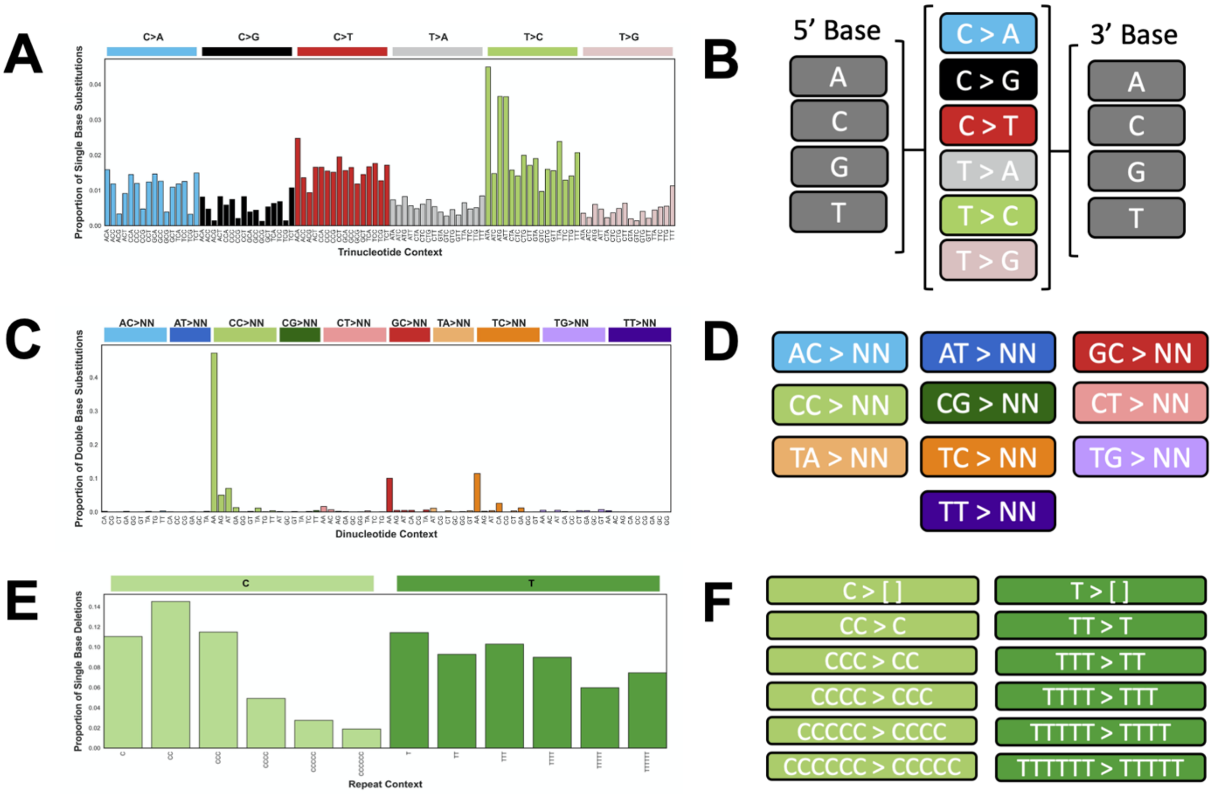


**Supplementary Figure S1**: **Mutational signatures classification scheme for single base substitutions, double base substitutions, and single base insertions and deletions**. **A.** Visualization of the proportion of 96 types of single base substitutions (SBS-96). **B.** SBS-96 mutational signatures are identified by the mutated pyrimidine base and the flanking 5' and 3' bases to the mutated base to form a trinucleotide context. **C.** Visualization of the proportion of 78 types of double base substitutions (DBS-78). **D.** DBS-78 mutational signatures are identified by 10 types of reference doublet bases mutated to an alternate doublet base. **E.** Visualization of the proportion of 12 types of single base deletions. 12 types of single base insertions are represented in the same scheme as the 12 types of single base deletions. **F.** Indel mutation types are identified by single pyrimidine base indels at mononucleotide repeat regions of lengths ranging from 1 base to 6 or more bases.

#### B SomaticSiMu Features

Simulated genomes with imposed known mutational signatures associated with cancer can be useful for evaluating the performance of machine learning-based classifiers of genomic sequences and mutational signature extraction tools. SomaticSiMu extracts known signature data from reference COSMIC mutational signatures and simulates SBS, DBS, and single base indels onto an input sequence based on known mutational signatures and user-specified parameters (see Figure S2 for standard workflow).

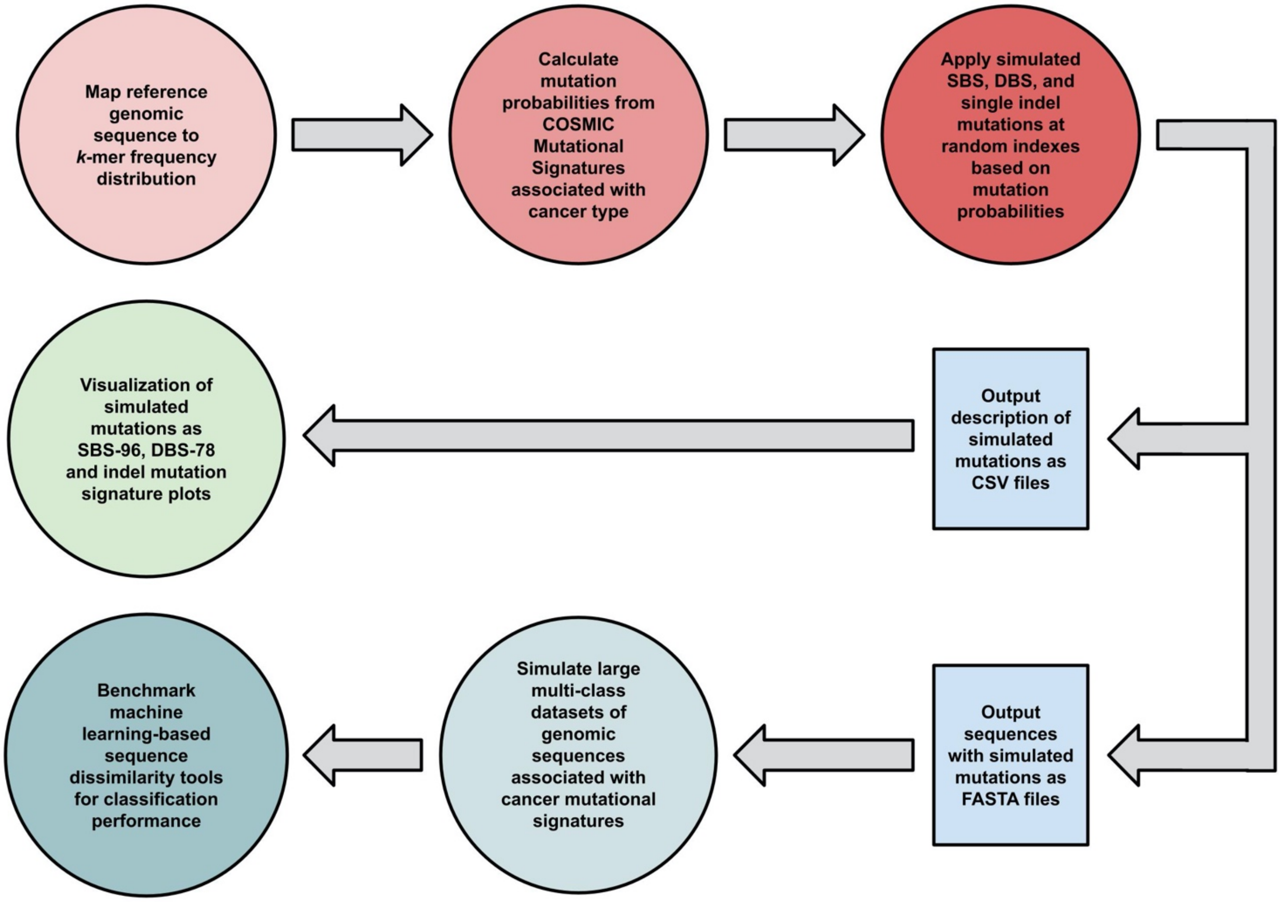


**Supplementary Figure S2**: **SomaticSiMu simulation workflow.** Simulated sequences and mutational catalogues are used to evaluate the accuracy of sequence classification of genomic sequences and the sensitivity of mutational signature extraction tools.

### C Simulation

#### C.1 Simulation Pipeline

1. *Index input sequence*

SomaticSiMu maps an indexed hash map of the index of every non-degenerate *k*-mer of 6 bases in length in the input sequence. The hash map is implemented as a Python dictionary object.

2. *Sampling signatures*

SomaticSiMu samples with replacement a subset of SBS, DBS, insertion and deletion mutational signatures based on the pervasiveness of each signature's activity in the user-specified cancer type from the 2,780 whole genome sequences in the reference Catalogue of Somatic Mutations in Cancer (COSMIC) Mutational Signatures v3.1 dataset. For example, mutational signatures that appear active in 50% of the variant sequences of one cancer type in the reference COSMIC dataset will be proportionally sampled and appear active in 50% of the simulated sequences. By default, the COSMIC dataset is filtered before signature sampling by excluding hyper-mutant sequences with a total sequence mutational burden above three standard deviations of the mean sequence mutational burden for each cancer type.

3. *Calculating signature mutational burdens*

Signature mutational burdens are defined as the total number of mutations attributed to one mutational signature active in one whole genome sequence. Mean signature mutational burden for each cancer was derived by summing the total number of mutations attributed to each signature in each tumor sample in the reference COSMIC dataset and dividing the sum by the number of sequences for each cancer. For example, the mean signature mutational burden for the SBS1 mutational signature active in the user-selected cancer type in the reference COSMIC dataset was calculated by summing the total mutational burden attributed to SBS1 across all 35 variants, then dividing by the number of tumor samples (n=35). This process was repeated to calculate the mean signature burden for all mutational signatures operative in the user-selected cancer type.

4. *Calculating mutation probabilities*

Mutation probabilities are defined as the probability that any given mutation in a local *k*-mer context will occur within one iteration of the simulation. SBS-96 mutational signatures are identified using the 3-mer trinucleotide context which includes the mutated base and the flanking 5' and 3' base. DBS-78 mutational signatures are identified using the 2-mer dinucleotide context which include the two consecutive mutated bases only. Single base insertion and single base deletion mutational signatures are identified using the 6-mer context.

Whole genome mutational burden based on the SBS-96 and DBS-78 mutational signatures as well as single base indel signatures for a user-specified cancer type are calculated by multiplying the mean signature mutational burden for each signature by the signature mutational proportions to output the mean number of each mutation type expected to appear in the human whole genome: Genome Reference Consortium Human Build 38 (GRCh38.p13; NCBI RefSeqAccession: GCF_000001405.26). For example, the whole genome SBS mutational burden of the user-selected cancer type in the reference COSMIC dataset based on the SBS-96 classification scheme was calculated by multiplying the mean signature burden for each active signature with the respective signature's 96-type mutational proportions to produce the mean whole genome mutational burden for each of the 96 mutation types. The mutational burden for each of the 96 mutation types across each of the active SBS signatures in the user-selected cancer type were summed together to produce the whole genome mutational burden using the 96 mutation type classification.

Mutation probabilities were calculated by dividing the whole genome mutational burden by the frequency distribution of *k*-mers in the GRCh38.p13 reference genome. For example, the SBS 96-type mutation probabilities of the user-selected cancer type were calculated by dividing the mutational burden of each of the 96 SBS mutation types by the total frequency of the respective 3-mer trinucleotide context in the GRCh38.p13 reference genome. If the input sequence is a subsequence of the Genome Reference Consortium Human Build 38 (GRCh38.p13; NCBI RefSeqAccession: GCF_000001405.26), mutation probabilities will be normalized such that the mutation types and burdens simulated using the input genomic sequence will be proportional compared to GRCh38.p13.

5. *Simulating mutations*

SomaticSiMu uses the mutation probabilities based on the SBS-96, DBS-78 and single base indel classification schemes to determine if any given base(s) in the input sequence will be mutated. The algorithm visits each base in the input sequence linearly starting at the first base and will either insert the SBS, DBS, and indel mutation or pass through without effect based on the mutation probability. The simulated SBS, DBS, and indel mutations are spiked into the input sequence linearly and the indexed hashmap of the sequence is updated with the new indices. Pseudo-random reseeding of each simulation generates unique and independent simulated mutations. SomaticSiMu is not limited to the size or length of the input reference sequence. SomaticSiMu has been tested with genomic fragments of 10,000 to 100 M base pairs in length.

6. *Output*

SomaticSiMu outputs a simulated sequence with active mutational signatures associated with the user-specified cancer type and metadata associated with the simulated mutation indexes, mutational burdens, and sets of signatures associated with each simulated sequence. The metadata of the simulated mutations are categorized based on the COSMIC SBS-96, DBS-78 and indel signature classification schemes. The logged metadata allows quantitative analyses of sequence total mutational burden and simulated mutation types.

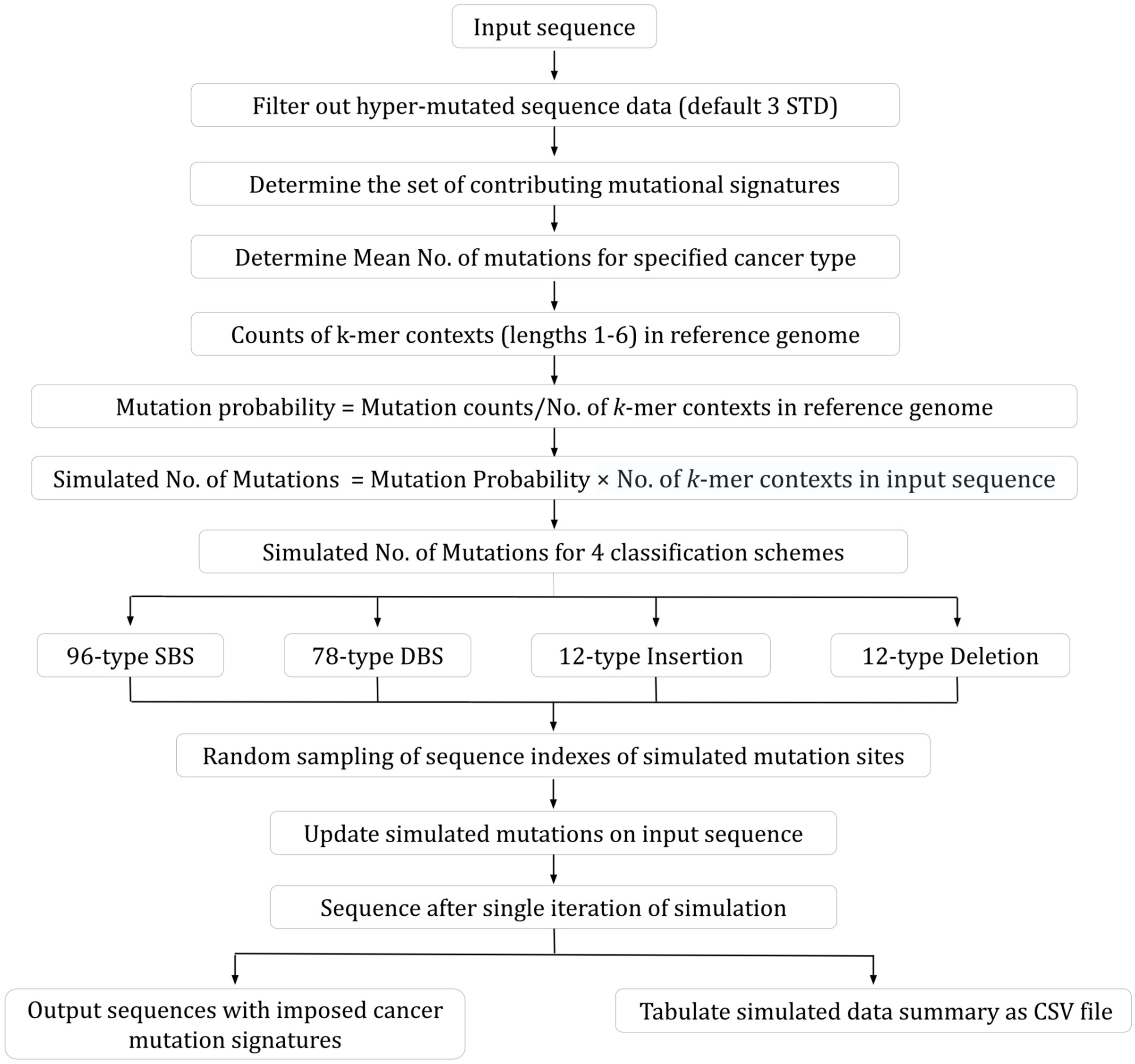


**Supplementary Figure S3: SomaticSiMu pipeline to generate simulated mutations by modelling mutational burdens and mutational signatures associated with human cancer.**

#### C.2 Simulation Parameters

The user can specify several parameters to calibrate their simulation experiment (Table S1).

**Supplementary Table S1**: **SomaticSiMu simulation parameters.**

| **Parameter** | **Short form parameter** | **Description** | **Default** |
|---|---|---|---|
| --generation | -g | Number of simulated sequences | 10 |
| --cancer | -c | Cancer type from the COSMIC dataset to simulate | |
| --reading_frame | -f | Start index of reading frame | 1 |
| --std | -s | Exclude signature data used to set up the simulation if sequence mutational burden is greater or less than n standard deviations from the mean | 3 |
| --slice_start | -a | Start index of the slice of the input sequence | None |
| --slice_end | -b | End index of the slice of the input sequence | None |
| --power | -p | Multiplier factor of mutational burden | 1 |
| --syn_rate | -x | Proportion of synonymous simulated mutations kept in the output simulated sequence | 1 |
| --non_syn_rate | -y | Proportion of non-synonymous simulated mutations kept in the output simulated sequence | 1 |
| --reference | -r | Absolute file path of reference sequence used as input for the simulation | |
| --normalization | -n | Normalize mutational probabilities proportional to Homo Sapiens Genome Assembly GRCh38.p13 whole genome. | False |

#### C.3 Simulation Output

SomaticSiMu names each simulated sequence file name and FASTA header with the simulated cancer type and one unique numerical identifier, starting at 1 and increasing by 1 for each sequence simulated. SomaticSiMu outputs the simulated sequences and associated metadata about the simulated sequences into 4 subdirectories (see Table S2).

**Supplementary Table S2**: **SomaticSiMu output directories.**

| **Directory Name** | **Description** | **Use Case** |
|---|---|---|
| Sample | Simulated sequences output into a subdirectory named after the type of cancer simulated within the Sample Directory. | Simulated FASTA sequences |
| Mutation_Metadata | CSV file output of each mutation simulated; the mutation type and index location on the reference input sequence. One file for each simulated sequence. | Identify type and exact location of each simulated mutation referenced by index of the input sequence for each simulated sequence. |
| Frequency_Table | CSV file output of summarized counts of each mutation type and local context. One file for each simulated sequence. | Sum of each simulated 96-type SBS mutation, 78-type DBS mutation, and 12-type single base indels for each simulated sequence. |
| Signature_Combinations | CSV file output of the signature combinations used for each iteration of the simulation. Different combinations of signatures are found operative in the same cancer type and are incorporated into the simulation. One file for each cancer type simulated. | Identify combinations of known SBS, DBS, and ID signatures used to model the simulated mutations for each simulated sequence with imposed cancer-associated mutational signatures. |

#### C.4 SomaticSiMu Graphical User Interface

SomaticSiMu is one of the first sequence simulators to simulate mutational signatures associated with human cancer and include a functional graphic user interface (GUI). The SomaticSiMu GUI features the same simulation capacity offered in SomaticSiMu, and allows users to intuitively set up their simulation experiments in a graphical environment (see Figure S4) as well as refer to the built-in graphical documentation by clicking on each input parameter without leaving the application. Moreover, the visualization menu of SomaticSiMu plots the metadata associated with the bootstrapped simulated sequences as SBS-96, DBS-78, single base indel mutational proportions and total sequence mutational burden plots.

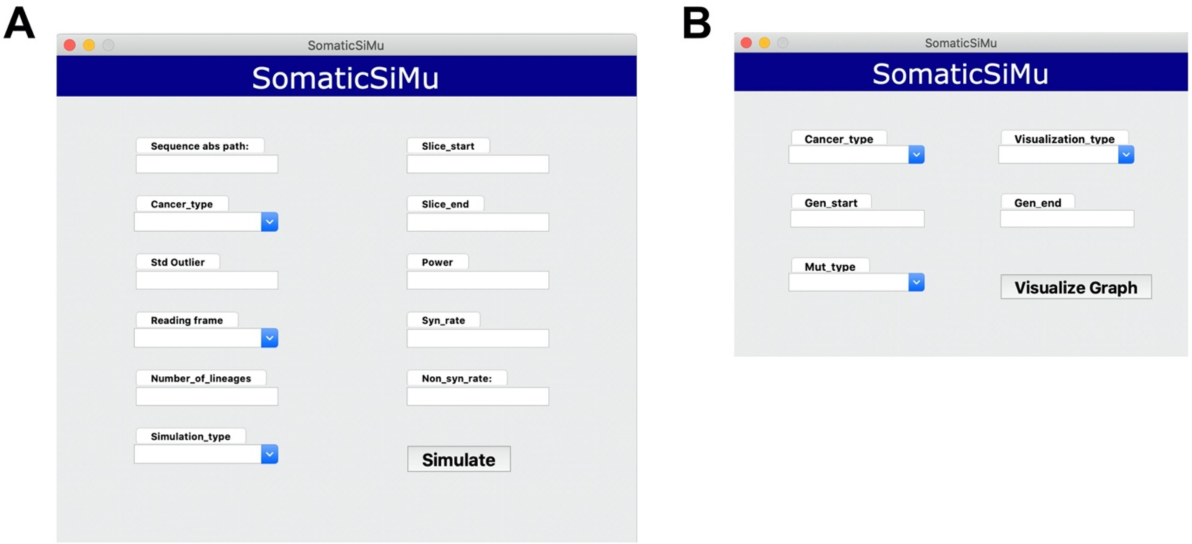


**Supplementary Figure S4: SomaticSiMu menu features a plug and play style interface with built-in graphical documentation. A.** SomaticSiMu Simulation Menu Parameters. **B.** SomaticSiMu Visualization Menu Parameters.

### D Visualization

SomaticSiMu features a set of visualization functions to plot the mean simulated SBS-96, DBS-78, single base insertion and deletion mutational proportions as well as mutational burdens using the output metadata in the Frequency_Table subdirectory. The plots visualize two simulated biomarkers: total sequence mutational burden and the catalogue of mutation types.

#### D.1 Single base substitution (SBS-96)

For SBSs, the predominant classification scheme used in SomaticSiMu includes 96 stand-agnostic mutation classes based on a single base substitution referred by its pyrimidine base as well as 5’ and 3’ bases adjacent to the mutated base that comprise the trinucleotide contexts as part of the SBS-96 classification, as visualized in Figure S5.

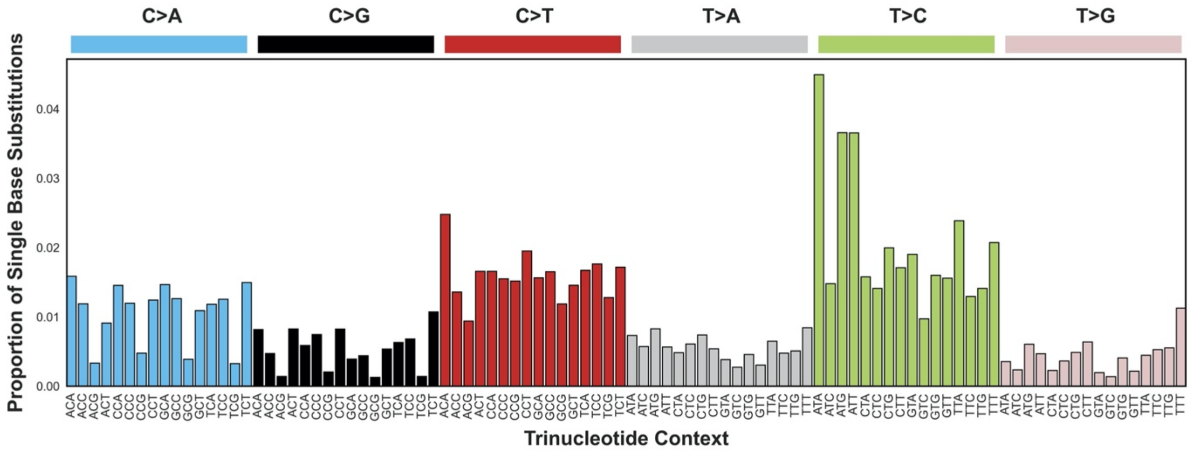


**Supplementary Figure S5**: **SomaticSiMu SBS-96 mutational proportions visualization.**

##### D.2 Double base substitution (DBS-78)

For DBSs, the predominant classification scheme used in SomaticSiMu includes 78 strand-agnostic mutation classes based on a pair of simultaneous adjacent substitutions mutated to another pair of two adjacent bases, as visualized in Figure S6.

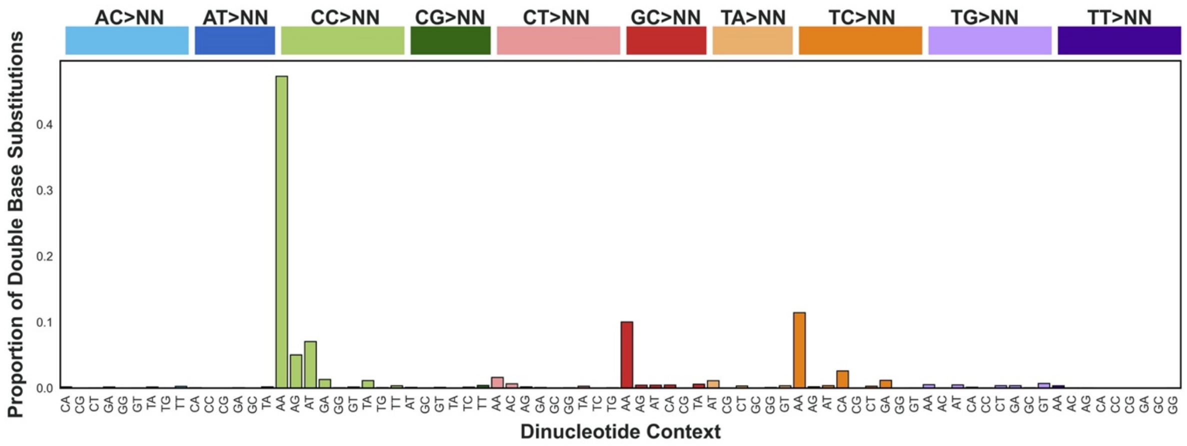


**Supplementary Figure S6**: **SomaticSiMu DBS-78 mutational proportions visualization.**

##### D.3 Single base insertion and deletion

For single base insertions and deletions, the classification scheme used in SomaticSiMu describes the indels by their occurrence at repetitive regions referred to as homopolymers, as visualized in Figure S7.

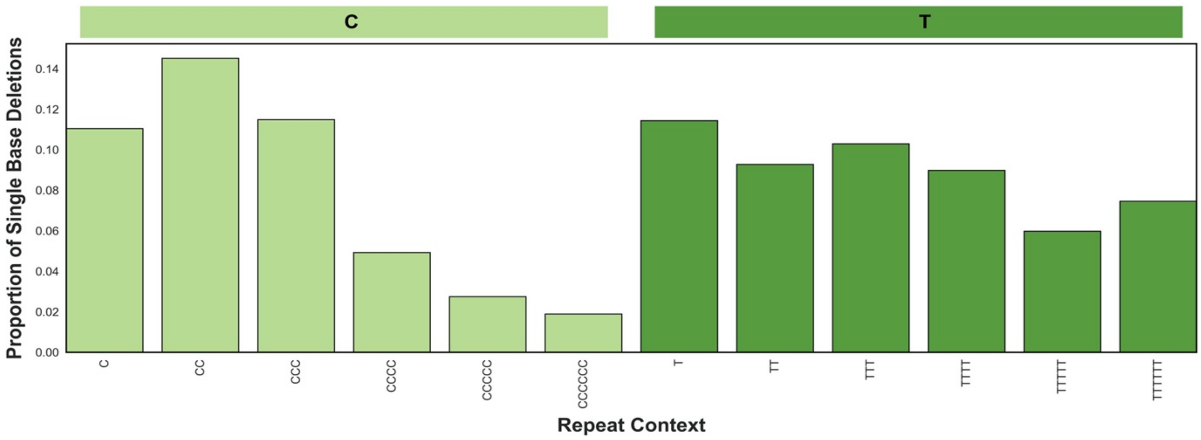


**Supplementary Figure S7**: **SomaticSiMu single base deletion mutational proportions visualization.**

##### D.4 Mutational Burden Plots

For total sequence mutational burden, the visualization function of SomaticSiMu plots the total number of simulated mutations for each sequence, categorized by mutation type. The mutation types simulated and visualized by SomaticSiMu includes SBS, DBS, and single base indels, as visualized in Figure S8.

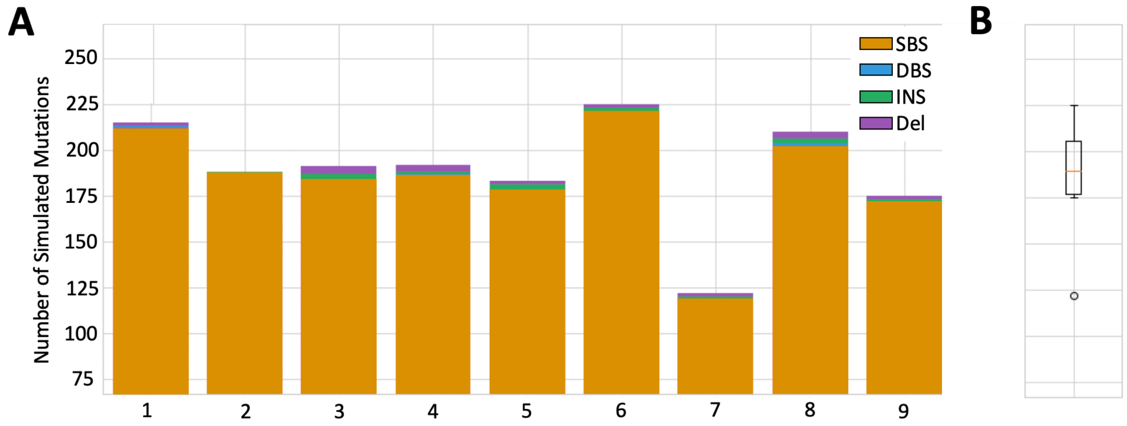


**Supplementary Figure S8**: **SomaticSiMu visualization of sequence mutation counts. A**. Stacked bar plots of the total number of SBS, DBS, and single base indel mutations for each unique simulated sequence. **B**. Distribution of total number of mutations of the all simulated sequences as a boxplot.

##### D.5 Visualization Parameters

The visualization function of SomaticSiMu requires that the CSV output associated with each simulation run exists in the Frequency_Table subdirectory of the SomaticSiMu directory. The user can specify several visualization parameters to set up visualizations depending on the use case (see Table S3). The user can use SomaticSiMu to plot the mean simulated SBS-96, DBS-78, and single base insertion and deletion mutational proportions as well as mutational burdens across multiple sequences at once by specifying the range of sequences using the Gen_start and Gen_end parameters. Gen_start and Gen_end specify the first sequence and last sequence respectively, and SomaticSiMu will visualize all sequences within the specified range, inclusive.

**Supplementary Table S3: SomaticSiMu simulation parameters.**

| **Parameter** | **Description** |
|---|---|
| Cancer_type | Cancer type to visualize |
| Visualization_type | Visualize using signatures associated with early or end (late) stage cancer |
| Gen_start | First simulated sequence metadata to visualize |
| Gen_end | Last simulated sequence metadata to visualize |
| Mut_type | Type of mutation proportions visualization |

### E Comparison between simulated data and real datasets

Simulated genomic sequences generated using SomaticSiMu mimic the same mutational types and proportions from the activity of multiple mutational signatures observed from 2,780 real whole cancer genomes from the Pan Cancer Analysis of Whole Genomes (PCAWG) Consortium 2020 study (Alexandrov *et al*., 2020). SomaticSiMu aims to simulate genomic sequences that have been impacted by multiple sources of somatic mutagenesis observed in human cancer, each with a characteristic mutational signature, that cumulatively alter sequence composition. For instance, ultraviolet light exposure is associated with SBS Signature 7 and commonly observed in Skin Melanoma genomes. Thus, the resulting mutational profile of Skin Melanoma in both real and SomaticSiMu-simulated genomic sequences is biased with the same high proportion of C>T substitutions seen in SBS Signature 7, see Supplementary Figure S11 for visualization of the mutational profiles.

The limitation of simulating mutational signatures *in silico* is that real cancer genomes have been impacted by natural selection for phenotypically advantageous mutations at specific genic locations, such as driver mutations that confer tumor growth advantages or deleterious mutations that knockout tumor suppressor activity (Stratton *et al.,* 2009). In contrast, SomaticSiMu simulates mutations in biologically representative types and proportions like real cancer genomes, but at random locations across the entirety of the input genomic sequence. Thereby, the functional impact of a subset of somatic mutations and the resulting selection for advantageous traits is not simulated using SomaticSiMu. Moreover, types of somatic mutagenesis observed in real cancer genomes includes but are not limited to the small-scale base substitutions and indels simulated by SomaticSiMu. For example, complex genomic alterations observed in real cancer genomes such as kataegis (Nik-Zainal *et al*., 2012) and chromothripsis (Cortés-Ciriano *et al*., 2020) are not simulated by SomaticSiMu and are outside the scope of the study at this time.

Benchmark tests were conducted to compare the similarity between the mutation types and proportions of real, ground-truth human cancer mutation catalogues sourced from COSMIC and cancer mutation catalogues simulated using SomaticSiMu. For each simulation, Human Chromosome 15 from GRCh.p38 (NCBI accession: NC_000015.10) was used as the input sequence and conducted simulations of 500 mutated sequences each for three cancer types using SomaticSiMu: Breast Adenocarcinoma (see Figure S9), Liver Hepatocellular Carcinoma (see Figure S10), and Skin Melanoma (see Figure S11). Visualizations of the mean proportion of each mutation type were plotted using the SBS-96, DBS-78, and single base insertion and deletion at repeat context classification schemes. Cosine correlation similarity ($cos(\theta)$), a measure of similarity between vectors, was used to evaluate the closeness between real and simulated data by comparing the mean vector of mutation types and proportions. Cosine similarity ranges from 0 to 1, where a similarity closer to 0 suggests dissimilarity between two vectors and a similarity closer to 1 suggests increased similarity between two vectors. Two-sided Wilcoxon rank-sum test is a test of difference between two paired groups used to test the null hypothesis that the mean mutational proportions of real and simulated mutations came from the same distribution. A p-value of less than 0.05 derived from the Wilcoxon rank-sum statistic indicates that the test rejects the null hypothesis at the 5% significance level.

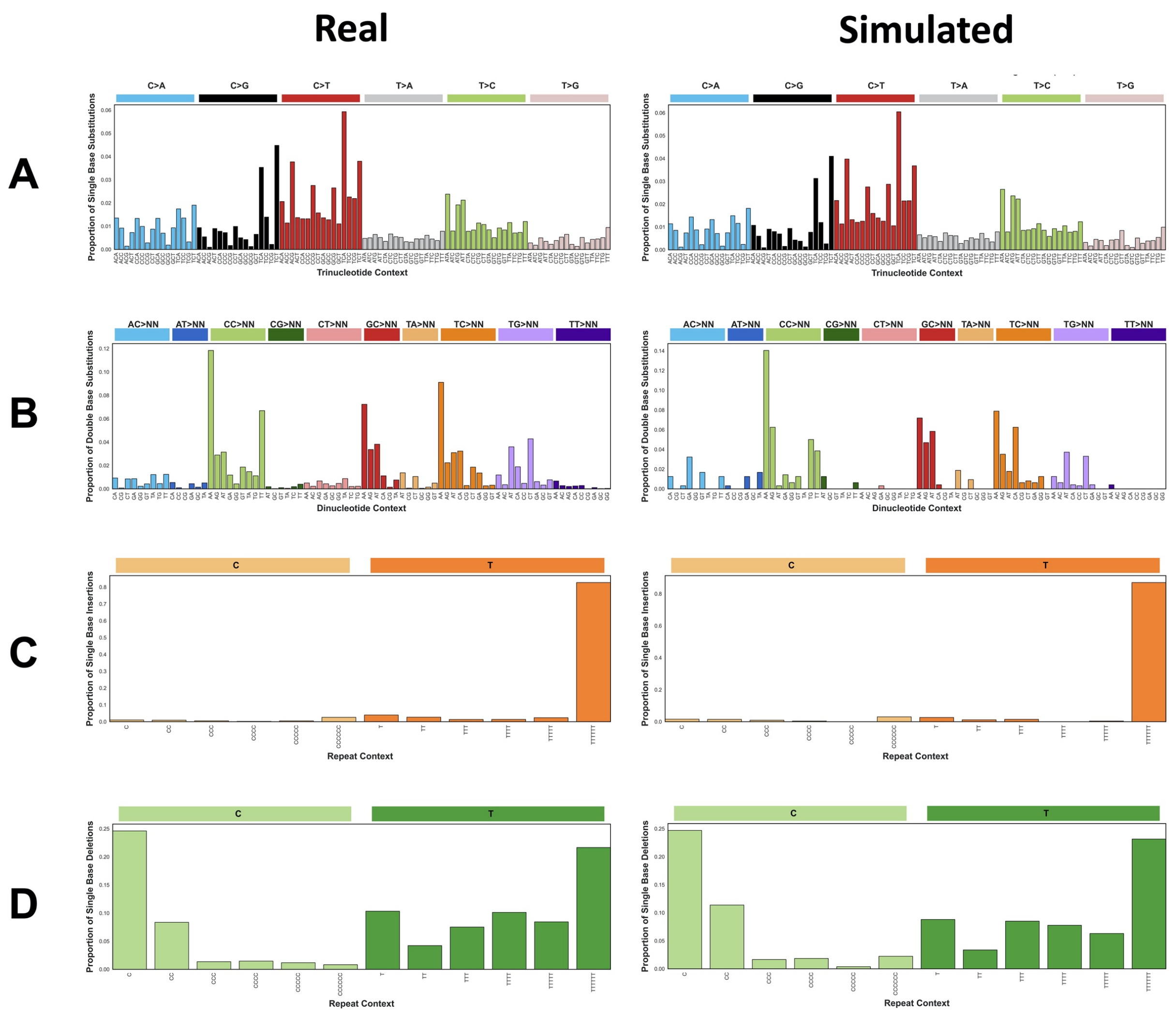


**Supplementary Figure S9**: **Comparative benchmark of the mean proportions of mutation types between real data (n=197) and simulated data (n=500) for Breast Adenocarcinoma. A.** SBS-96 plot of mutation types and proportions with reference to the mutation site trinucleotide context. High cosine similarity ($cos(\theta)$= 0.996) and no significant difference (p = 0.803, two-sided Wilcoxon rank-sum test) reported between real and simulated SBS mean mutational proportions. **B.** DBS-78 plot of mutation types and proportions with reference to the mutation site dinucleotide context. High cosine similarity ($cos(\theta)$= 0.958) and no significant difference (p = 0.409, two-sided Wilcoxon rank-sum test) reported between real and simulated DBS mutational proportions. **C.** Single base insertion plot of mutation types and proportions with reference to length of the mutation site repeat context. High cosine similarity ($cos(\theta)$= 0.999) and no significant difference (p = 0.729, two-sided Wilcoxon rank-sum test) reported between real and simulated single base insertion mutational proportions. **D.** Single base deletion plot of mutation types and proportions with reference to length of the mutation site repeat context. High cosine similarity ($cos(\theta)$= 0.994) and no significant difference (p = 0.862, two-sided Wilcoxon rank-sum test) reported between real and simulated single base deletion mutational proportions.

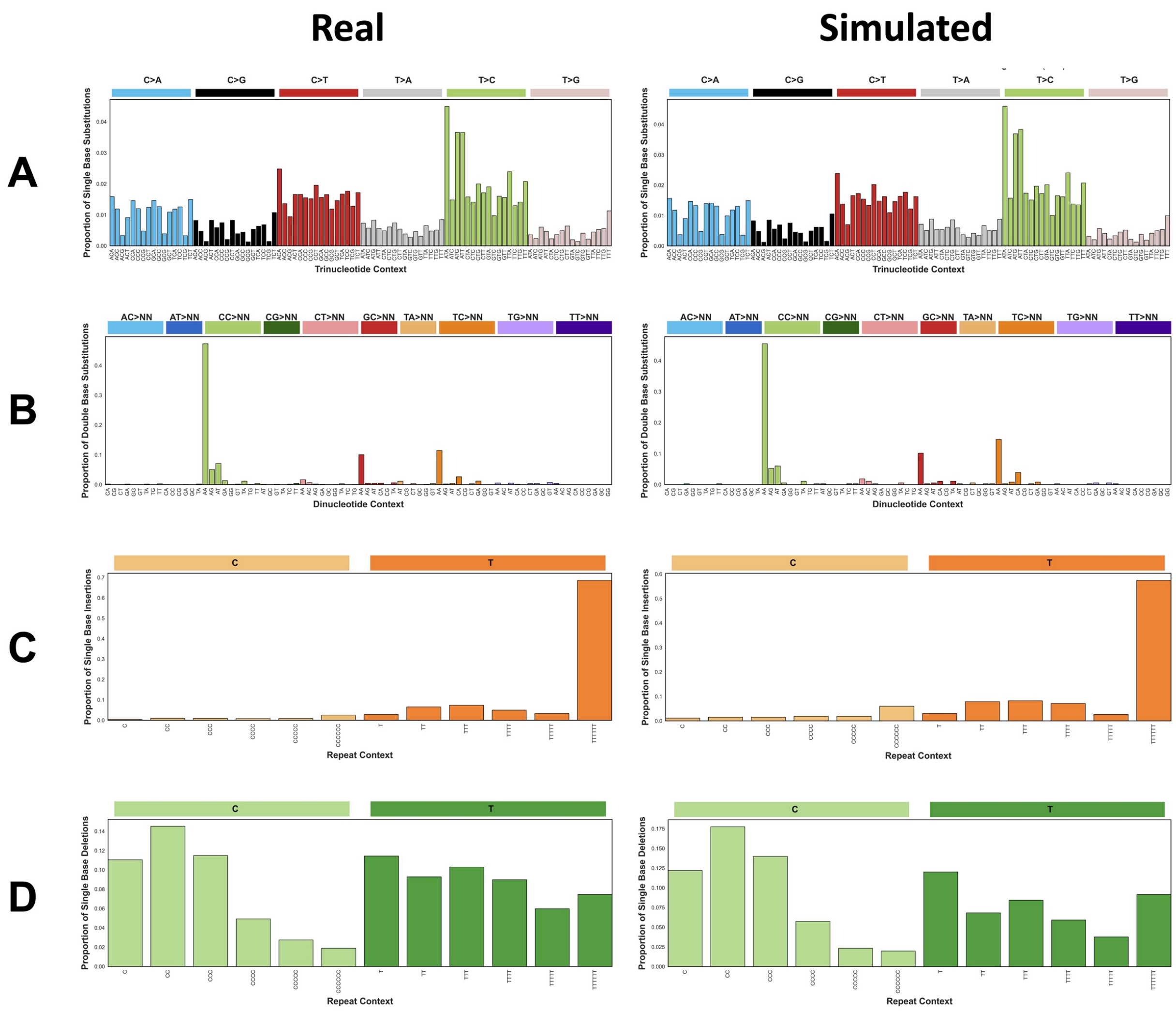


**Supplementary Figure S10: Comparative benchmark of the mean proportions of mutation types between real data (n=325) and simulated data (n=500) for Liver Hepatocellular Carcinoma. A.** SBS-96 plot of mutation types and proportions with reference to the mutation site trinucleotide context. High cosine similarity ($cos(\theta)$= 0.999) and no significant difference (p = 0.971, two-sided Wilcoxon rank-sum test) reported between real and simulated SBS mutational proportions. **B.** DBS-78 plot of mutation types and proportions with reference to the mutation site dinucleotide context. High cosine similarity ($cos(\theta)$= 0.998) and significant difference (p = 0.0244, two-sided Wilcoxon rank-sum test) reported between real and simulated DBS mutational proportions. **C.** Single base insertion plot of mutation types and proportions with reference to length of the mutation site repeat context. High cosine similarity ($cos(\theta)$= 0.996 ) and no significant difference (p = 0.356, two-sided Wilcoxon rank-sum test) reported between real and simulated single base insertion mutational proportions. **D.** Single base deletion plot of mutation types and proportions with reference to length of the mutation site repeat context. High cosine similarity ($cos(\theta)$= 0.996) and no significant difference (p = 0.954, two-sided Wilcoxon rank-sum test) reported between real and simulated single base deletion mutational proportions.

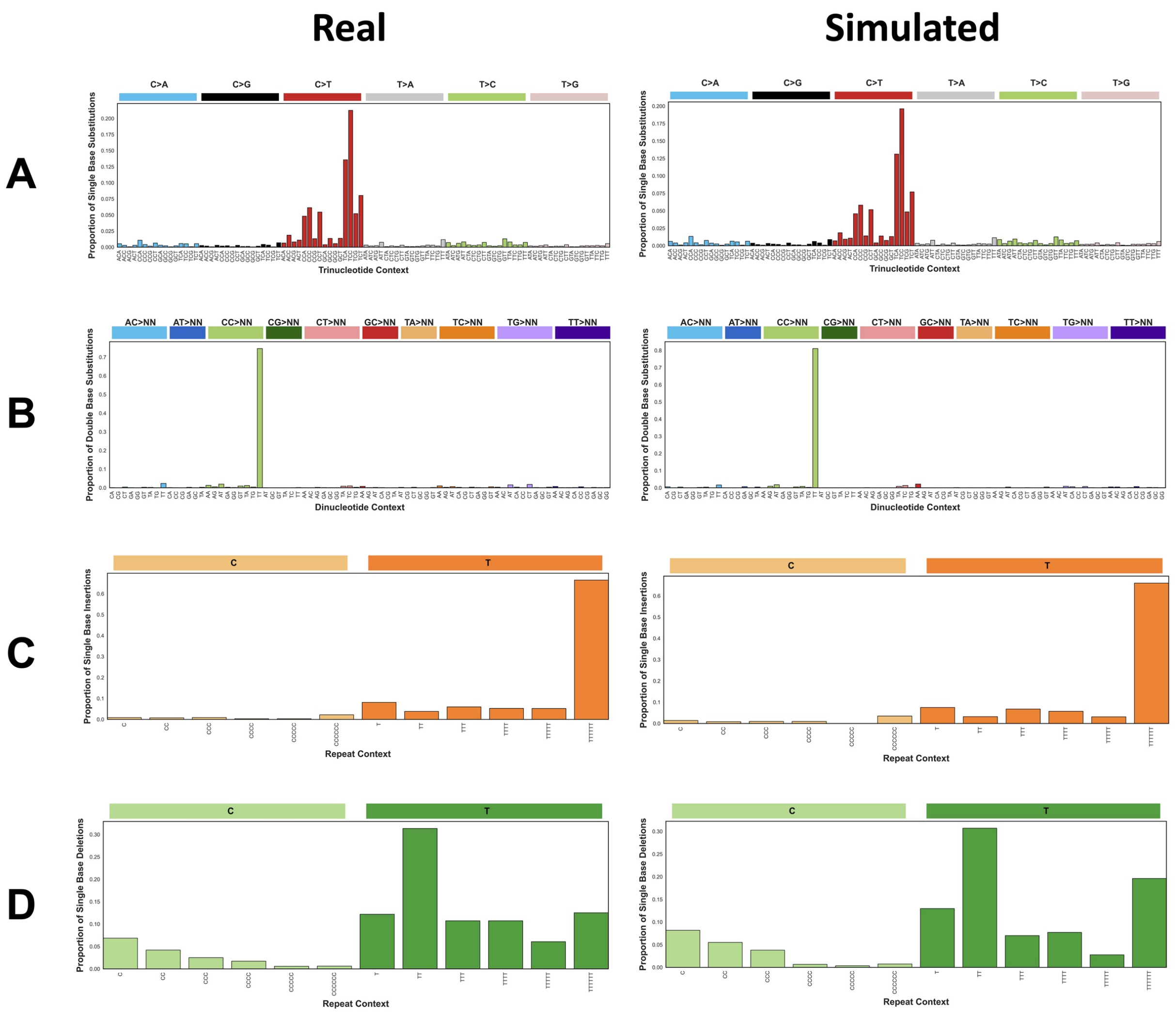


**Supplementary Figure S11**: **Comparative benchmark of the mean proportions of mutation types between real data (n=106) and simulated data (n=500) for Skin Melanoma.** Cosine similarity measured the correlation similarity between real and simulated mean proportions of mutation types. **A.** SBS-96 plot of mutation types and proportions with reference to the mutation site trinucleotide context. High cosine similarity ($cos(\theta)$= 0.999) and no significant difference (p = 0.860, two-sided Wilcoxon rank-sum test) reported between real and simulated SBS mutational proportions. **B.** DBS-78 plot of mutation types and proportions with reference to the mutation site dinucleotide context. High cosine similarity ($cos(\theta)$= 0.999) and no significant difference (p = 0.147, two-sided Wilcoxon rank-sum test) reported between real and simulated DBS mutational proportions. **C.** Single base insertion plot of mutation types and proportions with reference to length of the mutation site repeat context. High cosine similarity ($cos(\theta)$= 0.999) and no significant difference (p = 0.773, two-sided Wilcoxon rank-sum test) reported between real and simulated single base insertion mutational proportions. **D.** Single base deletion plot of mutation types and proportions with reference to length of the mutation site repeat context. High cosine similarity ($cos(\theta)$= 0.994) and no significant difference (p = 1.000 two-sided Wilcoxon rank-sum test) reported between real and simulated single base deletion mutational proportions.

### F Evaluating the reliability of simulated sequences using sequence classification tools

SomaticSiMu can simulate FASTA sequences with imposed mutational signatures associated with human cancer types. Machine Learning with Digital Signal Processing (MLDSP) is an alignment-free supervised machine learning tool used to classify FASTA sequences (Randhawa *et al*., 2019; Randhawa *et al*., 2020). MLDSP uses the simulated FASTA sequence datasets categorized by cancer type as input, preprocesses the DNA sequences into alignment-free feature vectors, and conducts 10-fold cross-validation to classify each sequence into cancer types using six machine learning classifiers. The aim of supervised machine learning classification is to correctly identify the class label of a subset of sequences when trained on other similar sequences with known labels.

The simulation mode of SomaticSiMu was used to simulate sequences with imposed mutation types and proportions that mimicked the COSMIC mutational signatures associated with human cancer. The simulation used Chromosome 22 from the Genome Reference Consortium Human Build 38 as the input reference sequence (NCBI accession: NC_000022.11). Five simulated sequence datasets with combinations of high or low total sequence mutational burden and high or low similarity between mutation types were generated using SomaticSiMu. The classification accuracy of MLDSP across each of the five simulated sequence datasets was compared to examine if SomaticSiMu can faithfully reproduce the diversity of mutation types and total sequence mutational burden that are expected to show higher or lower sequence classification accuracy.

**Supplementary Table S4**: **Simulated dataset manifest used to assess the reliability of SomaticSiMu and classification accuracy of MLDSP, a sequence classification tool.**

| Simulated Dataset ID | Simulated Cancer Type | Number of Sequences for each Cancer Type | Relative Total Sequence Mutational Burden | Relative Similarity of SBS-96 Mutational Proportions between Cancer Types |
|---|---|---|---|---|
| 1 | Bone Benign | 100 | Low<br>(<5 mutations/Mb) | High<br>(Cosine Similarity >0.85) |
| | CNS Pilocytic Astrocytoma | | | |
| | Myeloid Myelodysplastic Syndrome | | | |
| | Myeloid Myeloproliferative Neoplasm | | | |
| | Thyroid Adenocarcinoma | | | |
| 2 | Bone Osteosarcoma | 100 | Low<br>(<5 mutations/Mb) | Low<br>(Cosine Similarity <0.85) |
| | Breast Lobular Carcinoma | | | |
| | Cervix Adenocarcinoma | | | |
| | Kidney Renal Cell Carcinoma | | | |
| | Pancreas Adenocarcinoma | | | |
| 3 | Bladder Transitional Cell Carcinoma | 100 | High<br>(>5 mutations/Mb) | High<br>(Cosine Similarity >0.85) |
| | Breast Adenocarcinoma | | | |
| | Breast Lobular Carcinoma | | | |
| | Cervix Squamous Cell Carcinoma | | | |
| | Head Squamous Cell Carcinoma | | | |
| 4 | Colorectal Adenocarcinoma | 100 | High<br>(>5 mutations/Mb) | Low<br>(Cosine Similarity <0.85) |
| | Esophageal Adenocarcinoma | | | |
| | Lung Adenocarcinoma | | | |
| | Lung Squamous Cell Carcinoma | | | |
| | Skin Melanoma | | | |
| 5 | Colorectal Adenocarcinoma | 100 | Very high<br>(>50 mutations/Mb) | Low<br>(Cosine Similarity <0.85) |
| | Esophageal Adenocarcinoma | | | |
| | Lung Adenocarcinoma | | | |
| | Lung Squamous Cell Carcinoma | | | |
| | Skin Melanoma | | | |

#### F.1 Testing the reliability of simulated sequences to faithfully reproduce mutational signatures

Two genomic sequence biomarkers that MLDSP can use to differentiate between cancer types are mutational types and their proportions, and total sequence mutational burden. Mutation proportions are defined as the proportion of each mutation type out of all mutations observed in a simulated sequence. Ground-truth datasets of simulated sequences can be used to compare classification performance of MLDSP under scenarios where the sequences with imposed mutational signatures from different cancer types have relatively high or low total mutational burden and relatively high or low similarity of mutation types.

SomaticSiMu is designed to generate simulated sequences with imposed mutational signatures associated with human cancer that reliably reconstruct the mutation types and burdens of real cancer type mutational signatures. If SomaticSiMu produces reliable simulated sequences with imposed mutational signatures, higher classification accuracy is expected when MLDSP is tested using simulated sequences with a high mutational burden and low cosine similarity between the mutational types and proportions of different cancer types. Conversely, lower classification accuracy is expected when MLDSP is tested using simulated sequences with a low mutational burden and high cosine similarity between the mutational types and proportions of different cancer types.

From the MLDSP classification tests of five simulated sequence datasets, the classification accuracy of the best performing classifier, Linear SVM, and the average of the six MLDSP classifiers reported the lowest classification accuracy when tested on simulated sequences with more similar mutation types and lower total sequence mutational burden. As the similarity between mutation types decreases and the total sequence mutational burden increases, the classification accuracy of MLDSP increases as expected for reliable simulated sequences, see Supplementary Figure S12. The MLDSP sequence classification tests confirm that SomaticSiMu can simulate sequences with different mutation types and total sequence mutational burdens that accurately reproduce the expected classification accuracy results.

#### F.2 Use Case 1: Testing the classification accuracy of sequence classification tools

Evaluating the classification performance of genomic sequence-based supervised classification tools such as Machine Learning with Digital Signal Processing (MLDSP) requires large training and testing datasets of unique sequences with known sets of mutation types and frequencies. Since real sequence datasets show imbalanced class distributions and underrepresentation of unique sequences for rare cancer genomic sequences, a simulation approach was used to generate sequence datasets. Simulated sequences *in silico* are used to artificially upsample minority classes to obtain a more balanced class distribution recommended for supervised machine learning-based sequence classification (Leevy *et al*., 2018).

The classification accuracy of MLDSP was compared across five simulated sequence datasets, see Supplementary Figure S12 for cancer type classification results. MLDSP achieved the highest average accuracy of 43.2% when classifying sequences with high total sequence mutational burden and dissimilar mutational types and proportions. In comparison, MLDSP achieved the lowest average accuracy of 31.6% when trained and tested classifying sequences with high total sequence mutational burden and similar mutational types and proportions. Simulated sequence datasets with higher mutational burden (>5 mutations/Mb) reported higher classification accuracy compared to simulated sequence datasets with lowerer mutational burden (<5 mutations/Mb).

Across four out of five tests, the Linear SVM classification model ranked first based on 10-fold cross-validated classification accuracy compared to other classification models in the same test. Conversely, the Linear Discriminant classification model ranked last based on 10-fold cross-validated classification accuracy across three out of five tests compared to other classification models in the same test. Tests suggest that the Linear SVM classification model consistently achieves relatively higher classification accuracy of simulated sequences with imposed cancer mutational signatures.

The relatively low classification accuracy of simulated sequences by cancer type was hypothesized to be due to the low total mutational burden when simulating sequences using a relatively small input genomic sequence (50 Mb Chromosome 22). The classification accuracy reported by MLDSP is expected to increase when classifying larger simulated genomic sequences with higher total mutational burden.

To test this, we examined the classification accuracy of simulated hyper-mutant sequences with a 10-fold increase in mutational burden in Dataset 5 (>50 mutations/Mb) compared to Dataset 4 (>5 mutations/Mb). The same cancer types and numbers of sequences were simulated for Dataset 4 and Dataset 5. The increased 10-fold cross-validated classification accuracy of the highest accuracy classifier, Linear SVM (51.2% to 72.4%), and the average classification accuracy across all classifiers (43.2% to 64.3%) support the hypothesis that total sequence mutational burden may be positively associated with classification accuracy as expected in a reliable simulation.

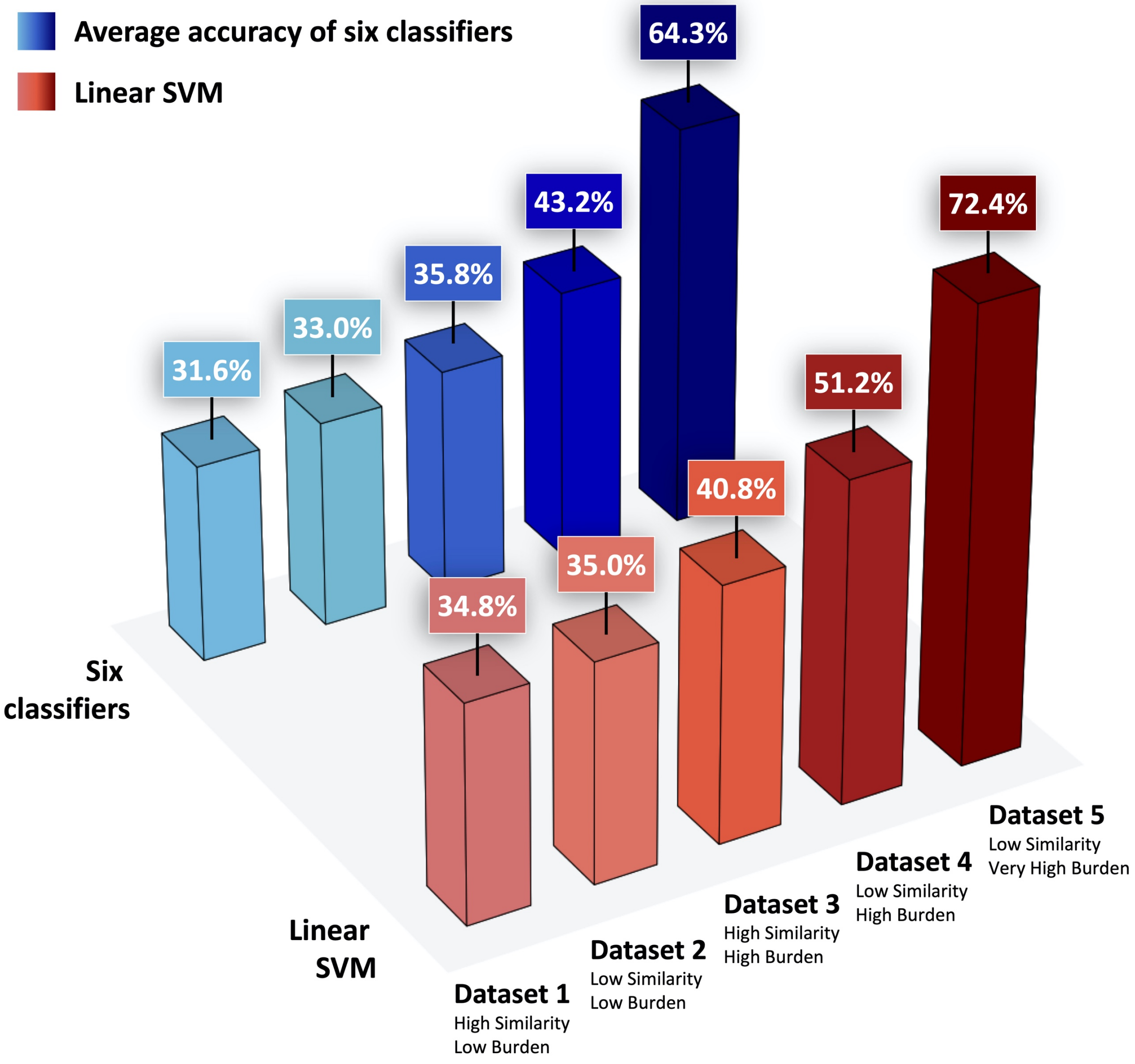


| MLDSP Classifiers | Dataset 1 Accuracy (High Similarity, Low Burden) | Dataset 2 Accuracy (Low Similarity, Low Burden) | Dataset 3 Accuracy (High Similarity, High Burden) | Dataset 4 Accuracy (Low Similarity, High Burden) | Dataset 5 Accuracy (Low Similarity, Very High Burden) |
|---|---|---|---|---|---|
| Linear Discriminant | 39.2% | 29.2% | 26.8% | 22.8% | 57.7% |
| Quadratic SVM | 35.8% | 36.4% | 34.6% | 37.4% | 58.3% |
| Fine KNN | 42% | 38% | 27.2% | 35.4% | 57.5% |
| Subspace Discriminant | 48% | 32.8% | 34.6% | 31.6% | 70.6% |
| Subspace KNN | 43% | 37.8% | 31.4% | 35.6% | 69.2% |
| **Linear SVM** | **51.2%** | **40.8%** | **34.8%** | **35%** | **72.4%** |
| **Average of 6 Classifiers** | **43.2%** | **35.8%** | **31.6%** | **33%** | **64.3%** |

**Supplementary Figure S12: MLDSP 10-fold cross-validated classification accuracy tested on five simulated sequence datasets outlined in Table S4.**

### G Evaluating the reliability of simulated mutational catalogues using mutational signature extraction tools

SomaticSiMu can generate count matrices, termed mutational catalogues, of simulated single base substitutions, double base substitutions, and single base indels associated with the imposed mutational signatures of each simulated genomic sequence. Each mutational catalogue describes the set of simulated mutations for one genomic sequence. Each simulation run samples a subset of the mutational signatures operative in the user-specified cancer type based on the COSMIC version 3.1 Mutational Signatures dataset. The output of simulated mutational catalogues and the set of simulated mutational signatures for each simulation run can be used as a ground truth dataset to evaluate signature extraction tools.

The simulation mode of SomaticSiMu was used to generate 100 simulated mutational catalogues with imposed mutation types and proportions that mimicked the COSMIC v3.1 mutational signatures associated with human Skin Melanoma. The simulation used Chromosome 15 from GRCh.p38 (NCBI accession: NC_000015.10) as the input reference sequence. The output of SomaticSiMu (100 simulated single base substitution mutational catalogues and 100 sets of simulated mutational signatures) was used to compare to signature fitting estimations from SigProfilerExtractor, a state-of-the-art tool for extraction of mutational signatures from catalogues of somatic mutations (Islam *et al*., 2021). Default parameters for SigProfilerExtractor were used, with the exception of setting the "opportunity_genome" parameter as "GRCh38". The signature extraction test using SigProfilerExtractor was conducted using 100 independent simulated mutational catalogues as input, where each catalogue represents the set of simulated mutation types and counts for each simulated sequence by SomaticSiMu.

##### G.1 Testing the reliability of simulated mutational catalogues to faithfully reproduce mutational signatures

Signature extraction tools that use a fitting approach aim to fit a mutational catalogue to a reference list of mutational signatures to estimate each reference signature's relative contribution to the overall mutational catalogue. Since the number and set of reference mutational signatures sampled for each simulated sequence is known as a ground truth, the sensitivity to which signature extraction tools can identify the set of simulated mutational signatures can be assessed. If SomaticSiMu generates reliable simulated mutational catalogues, signature extraction tools are expected to be able to consistently identify the same set of operative mutational signatures from simulated mutational catalogues with high mutational burdens. Conversely, a signature extraction tool would not be able to consistently identify the same set of operative mutational signatures from simulated mutational catalogues with low mutational burdens. Both sensitivity and specificity metrics range from 0% to 100%, with higher metrics indicating improved detection performance by the signature extraction tool.

Based on the extraction test, simulated mutational signatures with higher mutational burden reported higher sensitivity, such as SBS7a (79.5%) and SBS7b (71.1%). Conversely, simulated mutational signatures with lower mutational burden reported lower sensitivity, such as SBS7c (29.5%) and SBS7d (28.9%). The consistent extraction of mutational signatures with higher mutational burden and inconsistent extraction of mutational signatures with lower mutational burden as expected support the reliability of the simulated mutational catalogues generated by SomaticSiMu to realistically reproduce mutational signatures.

##### G.2 Use Case 2: Testing the sensitivity of mutational signature extraction tools

A true positive hit is defined as the event where SigProfilerExtractor correctly identified the same mutational signature used by SomaticSiMu to simulate the mutational catalogue. A true negative hit is defined as the event where SigProfilerExtractor correctly identified a mutational signature that was not used by SomaticSiMu to simulate the mutational catalogue. A false positive hit is defined as the event where SigProfilerExtractor incorrectly identified a mutational signature that was not used by SomaticSiMu to simulate the mutational catalogue. A false negative hit is defined as the event where SigProfilerExtractor incorrectly failed to identify a mutational signature that was used by SomaticSiMu to simulate the mutational catalogue.

The results of the signature extraction test using SigProfilerExtractor are shown in Figure S13. SigProfilerExtractor correctly identifies the dominant mutational signatures with high mutational burden observed in Skin Melanoma (SBS7a and SBS7b) in at least 70% of the simulated samples with the signature as true positive hits.

SigProfilerExtractor was highly sensitive in extracting SBS7a (79.5%) and SBS7b (71.1%). Moreover, SigProfilerExtractor was highly specific in failing to extract SBS2 (100%), SBS13 (100.0%), SBS17a (100.0%), SBS17b (98.0%), and SBS58 (100.0%) in the majority of the 100 simulated samples without the signature as true negative hits. Mutational signatures with lower mutational burden (SBS7c and SBS7d) failed to be extracted by SigProfilerExtractor in at least 55% of the simulated samples with the signature as false negative hits.

Further testing using different cancer types, user-specified parameters for SomaticSiMu, and user-specified parameters for signature extraction tools should be tested to identify the optimal use cases and limitations of different signature extraction tools.

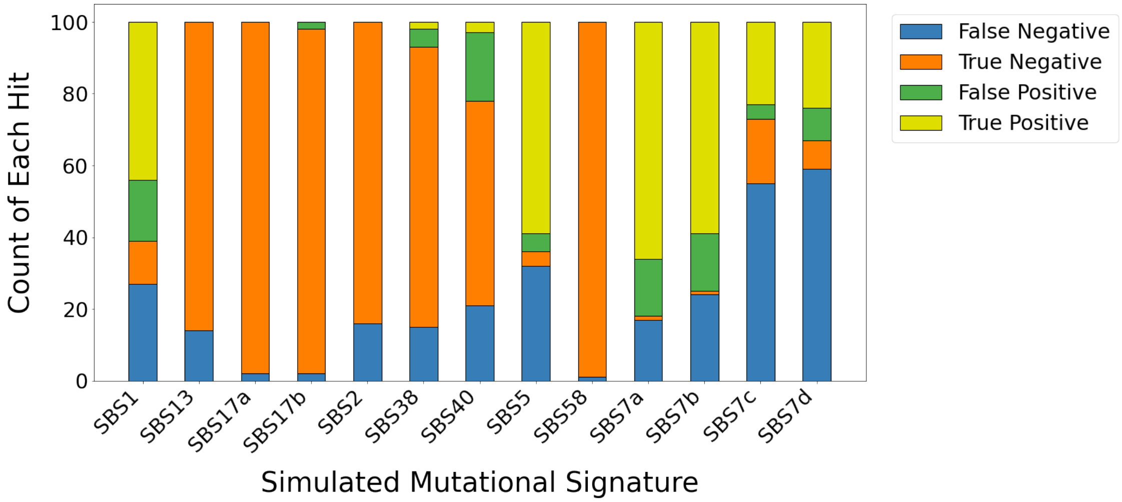


**Supplementary Figure S13**: **Stacked bar graph of the count of positive and negative hits of mutational signatures extracted from simulated mutational catalogues using the fitting approach by SigProfilerExtractor.** SomaticSiMu was used to simulate 100 unique mutational catalogues, each attributed to a set of mutational signatures associated with Skin Melanoma.

### H Installation

SomaticSiMu is hosted on GitHub and can be installed by cloning the SomaticSiMu repository from the GitHub repository to your local machine. The recommended instructions to clone the repository can be found in [GitHub Docs](). The setup script includes instructions to build, distribute, and install the minimum package dependencies needed for SomaticSiMu to run correctly. SomaticSiMu requires a minimal set of package dependencies which makes it deployable in both interactive use cases and production environments. The complete file structure expected after successful installation, including SomaticSiMu Python scripts, dependent reference datasets, and program metadata are listed below.

```
├── SomaticSiMu
│   ├── SomaticSiMu.py
│   ├── SomaticSiMu_GUI.py
│   ├── DBS_Expected_Frequency
│   ├── Documentation
│   ├── Frequency_Table
│   ├── kmer_ref_count
│   ├── ID_Expected_Frequency
│   ├── Mutation_Metadata
│   ├── Reference
│   ├── Reference_genome
│   ├── Sample
│   ├── Sample_Dataset
│   ├── SBS_Expected_Frequency
│   ├── Signature_Combinations
│   ├── kmer_ref_count
│   │   ├── 1-mer
│   │   ├── 2-mer
│   │   ├── 3-mer
│   │   ├── 4-mer
│   │   ├── 5-mer
│   │   ├── 6-mer
├── setup.py
├── README.md
├── LICENSE
├── requirements.txt
```

### I SomaticSiMu Quick Start Tutorial

The quick start tutorial provides two example scenarios for simulating genomic sequences with imposed mutational signatures and mutational burdens associated with a user-selected cancer type onto the entire length of input sequence Chromosome 22 from the Genome Reference Consortium Human Build 38 (NCBI accession: NC_000022.11). The same example simulation scenarios can be conducted using SomaticSiMu with a terminal interface and using SomaticSiMu with a graphical user interface as shown.

**Scenario 1**: Simulate 10 sequences by imposing known mutational signatures and mutational burdens associated with Biliary-Adenocarcinoma onto the entire length of input sequence Chromosome 22 (NCBI accession: NC_000022.11).

#### I.1 Scenario 1: SomaticSiMu command-line simulation

1. Run the SomaticSiMu simulation command provided below using a command-line interpreter. Update the sequence_abs_path parameter with the absolute file path of the sample input sequence Homo_sapiens.GRCh38.dna.chromosome.22.fasta found in the Reference_genome subdirectory included in the SomaticSiMu directory.

python SomaticSiMu.py --cancer Biliary-AdenoCA --reference
Homo_sapiens.GRCh38.dna.chromosome.22.fasta

2. The output simulated sequences and descriptive metadata of the successful simulation will be found in the Sample, Mutation_Metadata, Signature_Combinations, and Frequency_Table subdirectories in the SomaticSiMu directory.

#### I.2 Scenario 1: SomaticSiMu Graphical User Interface

1. Run the SomaticSiMu simulation command provided below using a command-line interpreter.

python SomaticSiMu_GUI.py

1. Input the SomaticSiMu simulation quick start parameters provided below into the SomaticSiMu simulation menu. Update the sequence_abs_path parameter with the absolute file path of the sample input sequence Homo_sapiens.GRCh38.dna.chromosome.22.fasta which is found in the Reference_genome subdirectory included in the SomaticSiMu directory.

sequence_abs_path = Homo_sapiens.GRCh38.dna.chromosome.22.fasta
cancer_type = Biliary-AdenoCA
std_outlier = 3
reading_frame = 1

number_of_lineages = 10
simulation_type = end
slice_start = 0
slice_end = 50818467
power = 1
syn_rate = 1
non_syn_rate = 1

2. Run *Simulate* to start the simulation.
3. The output simulated sequences and descriptive metadata of the successful simulation will be found in the Sample, Mutation_Metadata, Signature_Combinations, and Frequency_Table subdirectories in the SomaticSiMu directory.

##### I.3 Scenario 1: SomaticSiMu Graphical User Interface visualization

1. Input the SomaticSiMu visualization quick start parameters provided below into the SomaticSiMu visualization menu.

cancer_type = Biliary-AdenoCA
visualization_type = end
gen_start = 0
gen_end = 9
mut_type = SBS

2. Click *Visualize Graph* to begin the visualization
3. The visualization of the SBS mutations will be plotted in a new window. The window allows users to zoom in and out, resize, scroll, and save the plot.

**Scenario 2**: Simulate 100 sequences by imposing known mutational signatures associated with Breast Adenocarcinoma and increasing the mean mutational burden to twice as high as observed in real Breast Adenocarcinoma genomes onto the entire length of input sequence Chromosome 22 (NCBI accession: NC_000022.11).

##### I.4 Scenario 2: SomaticSiMu command-line simulation

3. Run the SomaticSiMu simulation command provided below using a command-line interpreter. Update the sequence_abs_path parameter with the absolute file path of the sample input sequence Homo_sapiens.GRCh38.dna.chromosome.22.fasta found in the Reference_genome subdirectory included in the SomaticSiMu directory.

python SomaticSiMu.py --cancer Breast-AdenoCA --reference Homo_sapiens.GRCh38.dna.chromosome.22.fasta --power 2 --generation 100

4. The output simulated sequences and descriptive metadata of the successful simulation will be found in the Sample, Mutation_Metadata, Signature_Combinations, and Frequency_Table subdirectories in the SomaticSiMu directory.

##### I.5 Scenario 2: SomaticSiMu Graphical User Interface

2. Run the SomaticSiMu simulation command provided below using a command-line interpreter.

```
python SomaticSiMu_GUI.py
```

4. Input the SomaticSiMu simulation quick start parameters provided below into the SomaticSiMu simulation menu. Update the sequence_abs_path parameter with the absolute file path of the sample input sequence Homo_sapiens.GRCh38.dna.chromosome.22.fasta which is found in the Reference_genome subdirectory included in the SomaticSiMu directory.

```
sequence_abs_path = Homo_sapiens.GRCh38.dna.chromosome.22.fasta
cancer_type = Breast-AdenoCA
std_outlier = 3
reading_frame = 1
number_of_lineages = 100
simulation_type = end
slice_start = 0
slice_end = 50818467
power = 2
syn_rate = 1
non_syn_rate = 1
```

5. Run *Simulate* to start the simulation.
6. The output simulated sequences and descriptive metadata of the successful simulation will be found in the Sample, Mutation_Metadata, Signature_Combinations, and Frequency_Table subdirectories in the SomaticSiMu directory.

##### I.6 Scenario 2: SomaticSiMu Graphical User Interface visualization

4. Input the SomaticSiMu visualization quick start parameters provided below into the SomaticSiMu visualization menu.

cancer_type = Breast-AdenoCA
visualization_type = end
gen_start = 0
gen_end = 99
mut_type = SBS

5. Click *Visualize Graph* to begin the visualization
6. The visualization of the SBS mutations will be plotted in a new window. The window allows users to zoom in and out, resize, scroll, and save the plot.

### J Availability

SomaticSiMu is open-source under the Creative Commons Attribution 4.0 International License terms. SomaticSiMu is available on GitHub at https://github.com/HillLab/SomaticSiMu.

### K Provided Datasets

SomaticSiMu provides four simulated datasets. All datasets were simulated using SomaticSiMu with Chromosome 22 from the Genome Reference Consortium Human Build 38 (NCBI accession: NC_000022.11) as the input sequence. Datasets are open source and can be downloaded at https://doi.org/10.5281/zenodo.5006275. Dataset details are given in Table S5. All simulated datasets were produced on a Macbook Pro A2141 using 8 cores of an Intel Core i9 9880H processor and 16GB DDR4 2667MHz SDRAM.

**Supplementary Table S5**: **Provided simulated datasets**

| ID | Dataset | Number of sequences | SomaticSiMu Simulation Time (seconds) |
|---|---|---|---|
| 1 | Colorectal Adenocarcinoma | 20 | 668 |
| 2 | Esophageal Adenocarcinoma | 20 | 155 |
| 3 | Lung Adenocarcinoma | 20 | 229 |
| 4 | Skin-Melanoma | 20 | 357 |
